# Decrease of Nibrin expression in chronic hypoxia is associated with hypoxia-induced chemoresistance in medulloblastoma cells

**DOI:** 10.1101/227207

**Authors:** Sophie Cowman, Yuen Ngan Fan, Barry Pizer, Violaine Sée

## Abstract

Solid tumours are less oxygenated than normal tissues. This is called tumour hypoxia and leads to resistance to radiotherapy and chemotherapy. The molecular mechanisms underlying such resistance have been investigated in a range of tumour types, including the adult brain tumours glioblastoma, yet little is known for paediatric brain tumours. Medulloblastoma (MB) is the most common malignant brain tumour in children. Here we used a common MB cell line (D283-MED), to investigate the mechanisms of chemo and radio-resistance in MB, comparing to another MB cell line (MEB-Med8A) and to a widely used glioblastoma cell line (U87MG). In D283-MED and U87MG, chronic hypoxia (5 days), but not acute hypoxia (24 h) induced resistance to etoposide and X-ray irradiation. This acquired resistance upon chronic hypoxia was much less pronounced in MEB-Med8A cells. Using a transcriptomic approach in D283-MED cells, we found a large transcriptional remodelling upon long term hypoxia, in particular the expression of a number of genes involved in detection and repair of double strand breaks (DSB) was altered. The levels of Nibrin (NBN) and MRE11, members of the MRN complex (MRE11/Rad50/NBN) responsible for DSB recognition, were significantly down-regulated. This was associated with a reduction of Ataxia Telangiectasia Mutated (ATM) activation by etoposide, indicating a profound dampening of the DNA damage signalling in hypoxic conditions. As a consequence, p53 activation by etoposide was reduced, and cell survival enhanced. Whilst U87MG shared the same dampened p53 activity, upon chemotherapeutic drug treatment in chronic hypoxic conditions, these cells used a different mechanism, independent of the DNA damage pathway. Together our results demonstrate a new mechanism explaining hypoxia-induced resistance involving the alteration of the response to DSB, but also highlight the cell type to cell type diversity and the necessity to take into account the differing tumour genetic make-up when considering re-sensitisation therapeutic protocols.

## Introduction

Medulloblastoma (MB) is a malignant embryonal brain tumour originating from neural stem cells or granule-cell progenitors of the cerebellum, due to a deregulation of signalling pathways involved in neuronal development such as Wnt or Sonic Hedgehog (SHH)^36^. It is one of the most common brain tumours in children accounting for 15-20% of all paediatric CNS tumours. MB is a heterogeneous cancer with distinct genetic variants, classified into 4 principal subgroups; Wnt, SSH, Group 3 and Group 4, based on their transcriptome^33, 63^ reviewed in^57^. More recent classification using genome wide DNA methylation and gene expression data resulted in the division of these 4 groups into 12 sub-groups^9^. Treatment of MB generally involves surgery followed by a combination of radiotherapy and cytotoxic chemotherapy. In very young children, surgery is followed by chemotherapy alone due to the severe effects of radiation therapy on the developing brain^48^. Recent studies have explored the use of proton beam therapy for MB, which triggers fewer detrimental side effects, as a result of reduction in irradiation of healthy brain tissue^68^.

Despite a marked improvement in the 5-year survival rate for MB patients, 40% succumb to the disease, demonstrating the limitations of the current therapies. Group 3 MB have the worst overall survival (~ 50%) even after extensive treatment^15, 43, 63^. One reason for treatment failure is the resistance to chemo- or radiotherapeutic interventions. In both MB patients and MB cell lines such resistance has been observed and attributed to mutations in specific signalling pathways in the case of targeted therapies^40^, or to mutations in the p53 signalling pathway^19, 23, 29, 39, 62^. For example, in the SHH group p53 mutations correlate with poor treatment outcome^70^. In addition to the intrinsic mutations in cancer cells conferring resistance, extrinsic factors such as tumour hypoxia have been implicated in chemo- and radioresistance in a range of tumour types^8 31, 47, 61^.

Tumour hypoxia is generated by irregular and tortuous vasculature formed within solid tumours, hence causing a poor delivery of oxygen and nutrients to cells. Hypoxia is associated with malignancy and tumour aggressiveness, by increasing tumour cell proliferation, metastasis and treatment resistance^27, 28, 55^. Hypoxic markers are, therefore, commonly associated with poor prognosis in many tumour types including breast cancer, glioblastoma and neuroblastoma^14, 24, 32, 65^. Interestingly, carbonic anhydrase IX (CaIX), a robust marker of hypoxic tumours^14^, is expressed in 23% of all MBs and is associated with poor prognosis^42^, suggesting that tumour hypoxia impacts on MB progression and/or management. The critical impact of tumour hypoxia on drug resistance and malignancy has been reported for a large number of cancers. However, very little is known about the effect of hypoxia on MB beyond, for example, a recent study showing that the hypoxia inducible factor (HIF-1) can influence the ability of MB cells to proliferate^18^.

Here we demonstrate that chronic hypoxia (>4 days) triggers resistance in a common MB cell line, D283-MED to cell death induced by etoposide or X-ray irradiation. The effect of chronic hypoxia on chemo- and radio-sensitivity was comparable to that observed in a classic glioblastoma cell line (U87MG). To understand the mechanisms underlying the observed resistance in the MB cells, we assessed changes in global gene expression driven by a longterm hypoxic exposure using a transcriptomic approach. We identified that the DNA double strand break (DSB) sensing machinery MRE11/RAD50/NBN, known as the MRN complex, was largely repressed. The MRN complex is recruited to sites of double-strand breaks as part of the Homologous Recombination Repair (HRR) and Non-Homologous End Joining (NHEJ) pathways. We further explored the consequences of such downregulation on the subsequent activation of the DNA damage response and apoptotic signalling pathways. Upon chronic hypoxia, we observed a reduced activation by etoposide of Ataxia Telangiectasia Mutated (ATM), a serine/threonine kinase recruited to double strand breaks. This was associated with reduced p53 stabilisation and transcriptional activity. Overall, this study demonstrates that the dampening of the DNA damage response to DSB triggered by chronic hypoxia strongly contributes to treatment resistance in hypoxic conditions in MB cells.

## Results

### Long term hypoxia induces etoposide and X-ray irradiation resistance in MB and glioblastoma cells

Hypoxia has previously been associated with increased drug resistance in a number of tumours including glioblastoma, hepatocellular carcinoma and breast tumours^1, 17, 61^. Medulloblastoma exhibit poor response to chemo- and radiotherapeutic interventions^25, 58^, yet the possibility that this could be explained by tumour hypoxia is unexplored. We used two MB cell lines: D283-MED (representative of Group3/4)^4, 54^, and MEB-Med8A (representative of Group 3)^35, 44^, to assess the effects of hypoxia pre-conditioning on cell sensitivity to chemotherapeutic treatment and X-ray irradiation. The results were compared with the classic glioblastoma cell line U87MG. Glioblastomas have been extensively described to be hypoxic, with oxygen concentration measured around 1% O_2_ or below^16, 66^, with a strong correlation between regional hypoxia and poor patient survival^60^ Here we used 1% O_2_ for hypoxic conditions as this level can induce a hypoxic response (increased levels of HIF-1α) without affecting cell proliferation or survival in a range of brain tumour cell lines^50^.

The D283-MED and MEB-Med8A cell lines display differential sensitivity to etoposide^23, 41^, hence a range of etoposide doses have been used. The cells were exposed to 1% O_2_ for 1 day (acute hypoxia) or for 5 days (chronic hypoxia) prior to etoposide treatment. The cells subsequently remained under hypoxic conditions during treatment. Control samples were cultured and treated at atmospheric O_2_ levels (normoxia, 21% O_2_). For D283-MED cells, MTS survival assays showed that 5 days of hypoxic preconditioning triggered significant resistance to etoposide, with a cell viability of ~74% in hypoxic cells treated with etoposide compared to ~14% in normoxic conditions (Figure 1A). In contrast, 1 day of hypoxic preconditioning did not induce any significant resistance, suggesting that the duration of hypoxic exposure is critical to alter the cell response to drugs. The hypoxia-induced resistance to etoposide was also observed in MEB-Med8A cells, although to a lesser extent and with high variability thereby without reaching statistical significance (Figure 1B; p=0.57). The U87MG glioblastoma cells displayed similar results to the D283-MED cell line (Figure 1C), with 5 day preconditioning resulting in significantly higher cell viability compared to the normoxic controls. Results for the D283-MED cell line were reproduced with the cell viability assay ViaCount™, to ensure that the MTS results were not biased by altered mitochondrial activity in hypoxia, and similar data were obtained (Figure S1A). Acquired resistance was also investigated upon X-ray irradiation. In this case, the cells were pre-incubated in 1% or 21% O_2_ for 5 days and the cells were irradiated in normoxic conditions to prevent loss of the oxygen enhancement effect during irradiation treatment^12^ With this protocol we ensured that only the effects of hypoxia preconditioning were assessed. Similarly to etoposide treatment, yet to a lesser extent, resistance to X-ray irradiation upon hypoxic preconditioning was observed for the D283-MED and the U87MG cell lines (Figure 1D). The MEB-Med8A cells showed an inverse trend (Figure 1D). This counterintuitive result may be due to the fact that this cell line appeared stressed by the cooling procedure prior to irradiation, hence generating variable results.

**Figure 1:**
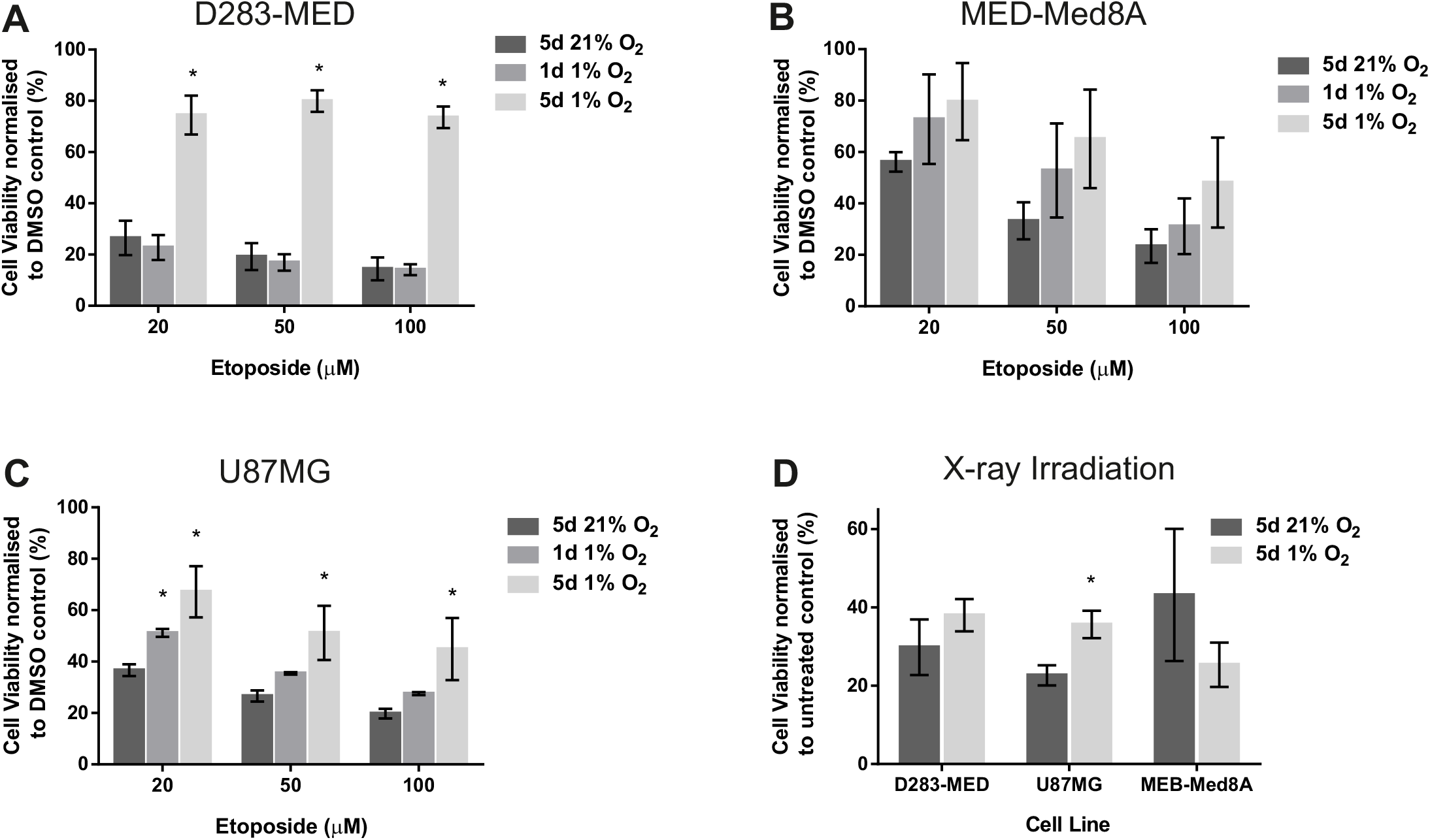
Chronic hypoxia-induces treatment resistance in MB and glioblastoma cells. Two MB and one glioblastoma cell lines were pre-cultured in normoxia (21% O_2_) or hypoxia (1% O_2_) for indicated time points prior to etoposide or X-ray irradiation treatment. Cells were maintained in 1% O_2_ during etoposide treatment [20-100 μM]. (A) D283-MED cells (B) MEB-Med8A (C) U87MG cells. (D) X-ray irradiation treatment was conducted in 21% O_2_ with doses of 30 Gy (D283-MED), 50 Gy (MEB-Med8A) and 2x80 Gy dose (U87MG), with 48 h incubation post-treatment. The different doses is to account for different cell sensitivity to irradiation. Cell viability was determined using an MTS assay, with absorbance normalised to untreated control. Data are shown as the mean ± S.E.M of at least 3 independent experiments. A student t-test was performed where (*) indicates statistical significance with p<0.05.

We further assessed the ability of etoposide to induce cell cycle arrest in D283-MED cells preconditioned or not in 1% O_2_. Hypoxia preconditioning on its own did not induce any cell cycle alteration (Figure S1B and C). This important control indicates that the reduced effect of etoposide in hypoxic preconditioned cells is not due to changes in cell proliferation agreeing with previous observations in glioblastoma^50^ Etoposide induced a clear G2/M arrest in normoxic cells (63% of cells in G2/M when treated with etoposide, compared to 30% for untreated cells). This arrest was largely diminished in hypoxic cells with only 40% in G2/M upon etoposide treatment (Figure S1B), in line with the effects of hypoxia on etoposide-induced cell death. Taken together, chronic hypoxia exposure results in resistance to the chemotherapeutic agent etoposide induced cell death and cell cycle arrest, and it also affects the sensitivity to X-ray irradiation. This resistance is not due to alteration of the cell cycle by hypoxia (Figure S1) or to the absence of oxygen during the irradiation process, as irradiation was performed in atmospheric conditions. It, therefore, points to global changes in cell signalling, resulting in lack of sensitivity to the DNA damaging protocols used.

### Chronic hypoxic exposure induces large transcriptional remodelling

To investigate the molecular mechanisms driving the observed hypoxia-induced cell death resistance in D283-MED, we used micro-array gene expression analysis to assess the global transcriptomic modifications triggered by long-term hypoxia. Hypoxia-inducible factors (HIF) are the main transcription factor family directly regulating gene expression in hypoxia. HIF-1 alpha (HIF-1α) is the primary isoform governing the expression of genes whose levels are directly dependent on oxygen availability. Hypoxia-induced gene expression was measured at a time point that correlates with the first peak in HIF-1α stabilisation, and at two later time points subsequent to additional peaks of HIF accumulation (Figure 2A). Considering all three time points together, we found that 6,124 transcripts were significantly up-or down-regulated in hypoxia (Figure 2B). This corresponds to 4,303 significantly regulated genes (and 23,611 non-regulated genes). Significantly regulated transcripts were further analysed globally by clustergram (Figure 2C), which showed a large number of genes regulated at late time points (64 h and 96 h), but not in acute conditions (6 h) (Figure 2D, E). Most transcriptional analyses upon hypoxic exposure have previously been performed after 24-48 h of hypoxia, corresponding to more acute conditions. Given the high number of transcripts regulated only at 64 h and 96 h, it is likely that many regulatory mechanisms in cellular adaptation to long term hypoxia have been missed thus far.

**Figure 2:**
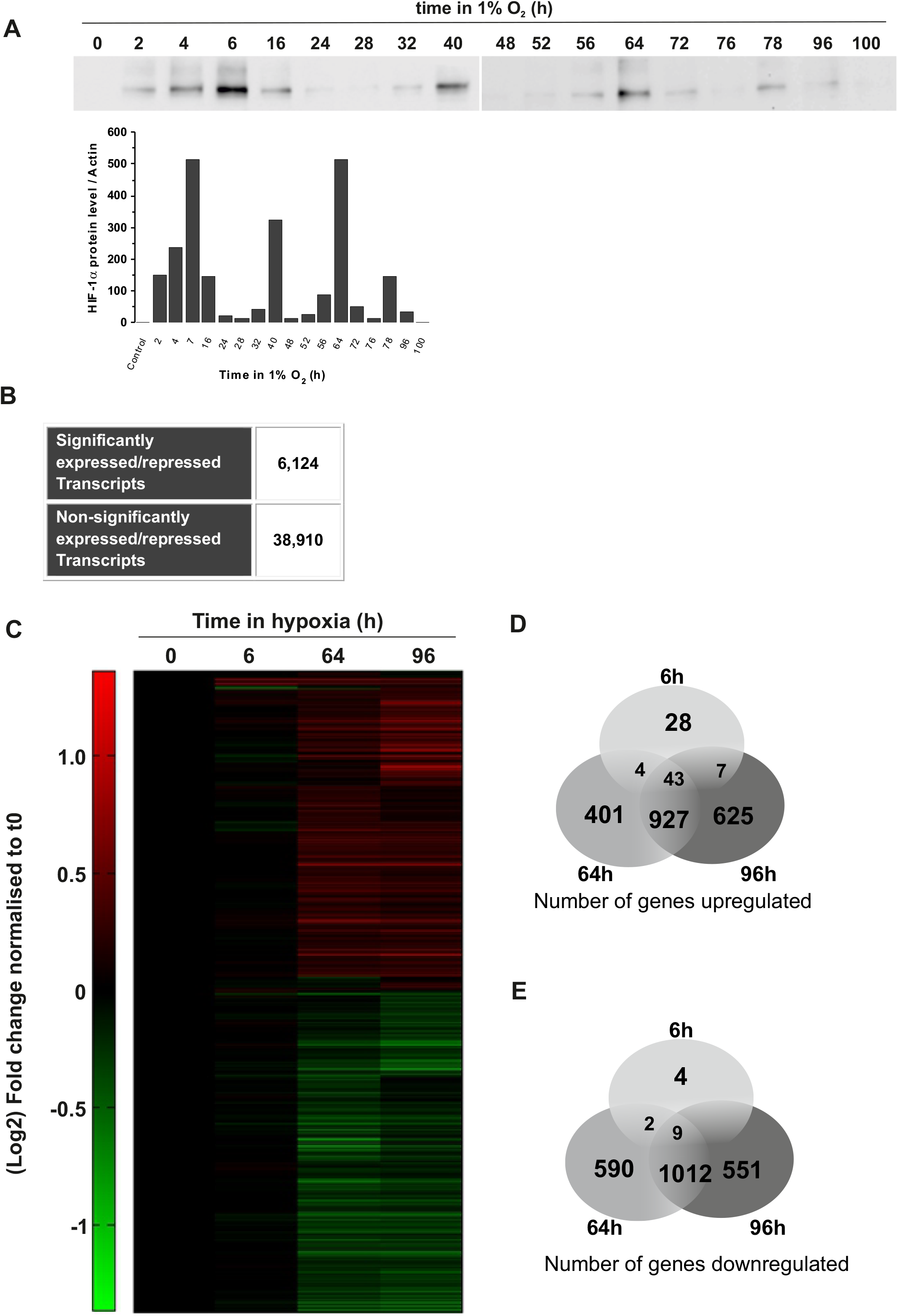
Hypoxia induces transcriptional remodelling. (A) D283-MED cells were incubated in 1% O_2_ for 0-100 h or 21% O_2_ as a control. The induction HIF-1α levels were measured by western blot. Data shown are representative of 2 independent experiments. (B) Number of significantly expressed or repressed transcripts in hypoxic time points, where significance represents transcripts which have either p<0.01 for all 3 probes, or p<0.0001 for 2 out of 3 probes, as well as a minimum of 2-fold change compared to normoxic control. (C) Clustered heat map representing the gene expression profile of D283-MED cells in hypoxia, where cells were incubated for 0, 6, 64 and 96 h at 1% O_2_. Green and red represent downregulation and upregulation respectively. Venn diagram depicting the number of (D) upregulated and (E) downregulated transcripts in D283-MED at each hypoxic time point.

We initially performed a biased analysis and specifically examined the expression of several multi-drug resistance genes. These genes were the obvious candidates potentially responsible for hypoxia-induced treatment resistance and had previously been described to have a role in hypoxia-induced drug resistance in other cellular models including glioblastoma^13^. However, none of the well described multi-drug resistance (MDR) genes *abcb1 (mdr1), abcc1 (mrp1)* and *abcg1* were up-regulated in hypoxia (Figure 3A).

**Figure 3:**
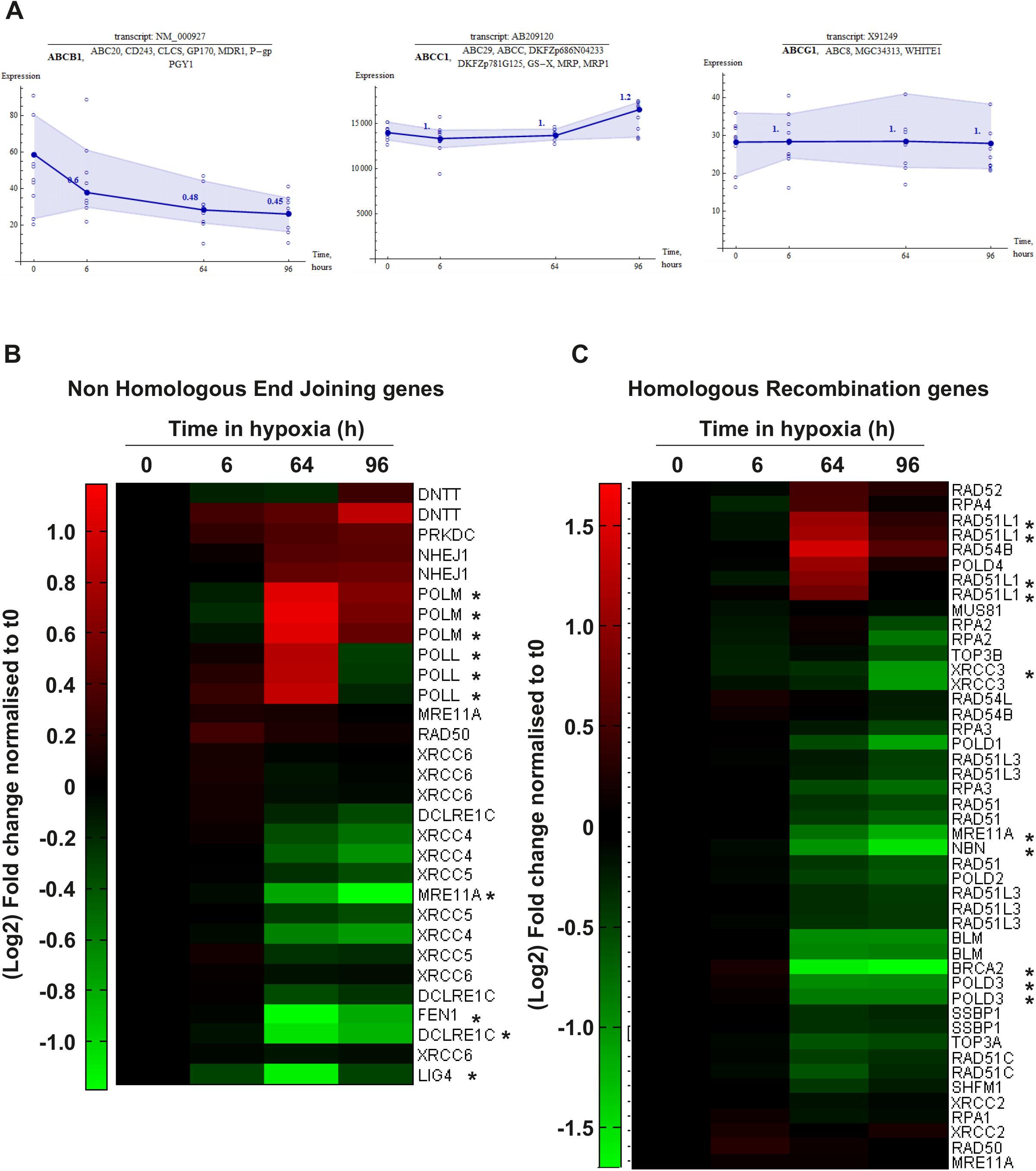
Hypoxia regulates DNA repair genes at the transcript level. (A) Expression plots for multidrug resistance genes, ABCB1, ABCC1 and ABCG1 in D283-MED cells incubated in 1% O2 for 0-96 h. Filled circle and dark blue line show the average expression of the transcript. Open circle represents the data point for each individual probe. Numbers above data points show the fold change normalised to basal ratio. Shaded blue areas indicates the range of data variation. Expression profiles of genes involved in (B) non-homologous end joining or (C) homologous recombination for D283-MED cells incubated at 1% O_2_ for 0-96 h. Green and red represent downregulation and upregulation respectively.

For a non-biased analysis using the *k*-means clustering method, the large number of regulated transcripts were further grouped into 16 smaller clusters based on their expression profile pattern, to find potential correlation between groups of genes with similar expression dynamics (Figure S2A). From this we identified that genes involved in the response to double strand breaks were strongly regulated at later hypoxic time points. The effectiveness of DNA damage response mechanisms can play a key role in response to chemo- and radiotherapy. Therefore, we generated heat map clustergrams of the HRR and NHEJ pathways, which are primarily responsible for the repair of DSB, the most lethal form of DNA damage (Figure 3B and C). Strong regulation was observed for a number of genes involved in both NHEJ (Figure 3B) and HRR (Figure 3C). We further focused on HRR pathways as key genes involved in signal transduction for HRR were down-regulated strongly by hypoxia. Down-regulated transcripts include *brca2* and two members of the MRN complex, *mre11A* (MRE11) and *Nibrin* (NBN). NBN plays a key role in ATM activation and signal amplification of the DNA damage response^10^ Therefore, the hypoxia mediated down-regulation of key components of the HRR pathway could be responsible of the altered cellular response to etoposide.

### DNA damage recognition and signal transduction is dampened in chronic hypoxia

Sensing of DNA damage is a crucial step in the initiation of double strand break (DSB) repair by the HRR machinery, and its alterations is likely to impact further downstream signalling of the repair pathway. The strong reduction in NBN transcription (70% decrease) (Figure 4A) observed in the micro-array was confirmed using qPCR, which equally showed a decrease of NBN mRNA by 70% at 96 hours (Figure 4B). Similarly, NBN protein levels were reduced by ~50% after 96 hours of hypoxic exposure (Figure 4C). To further understand the effect of chronic hypoxia on each member of the MRN complex, mRNA levels of NBN, MRE11 and RAD50 were determined by qPCR upon 5 days of hypoxic incubation. Hypoxia had little effect on RAD50 mRNA levels, yet there was significant down-regulation of MRE11 (~53%) comparable to NBN (Figure 4D), which was further confirmed at the protein level (Figure 4E). Surprisingly, such decreases in NBN and MRE11 expression was not observed in the U87MG cells (Figure S3A).

**Figure 4:**
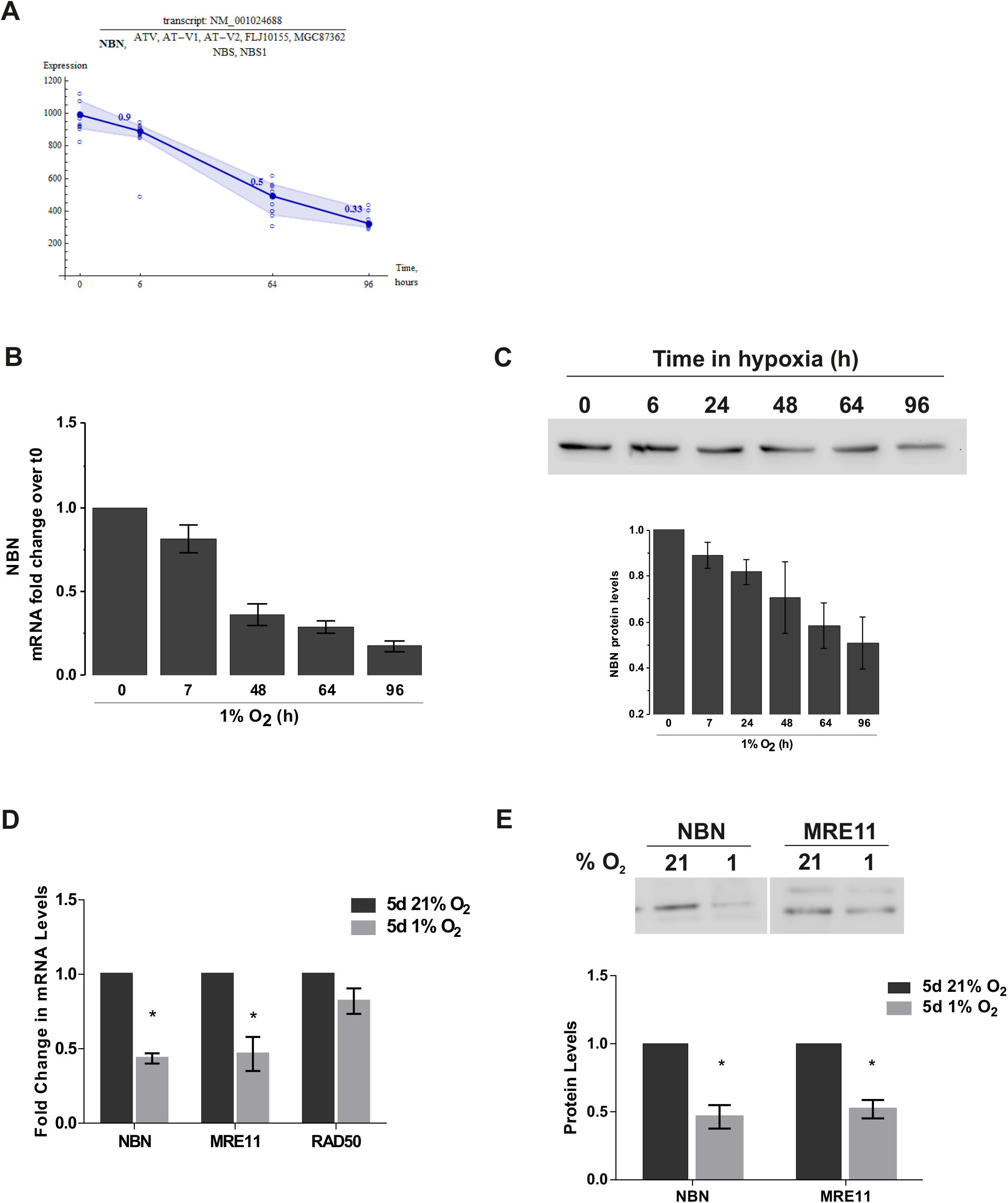
Reduction of Nibrin expression induced by chronic hypoxia. (A) Expression plots of NBN mRNA in D283-MED cells incubated in 1% O_2_ for 0-96 h. Filled circle and dark blue line show the average expression of the transcript. Open circle represents the data point for each individual probe. Numbers above data points show the fold change normalised to basal ratio. (B) NBN mRNA measured by qPCR normalised to cyclophillin A and the normoxic (21% O_2_) control. (C) NBN protein levels over time in hypoxia (0-96 h), measured by western-blot. (D) Levels of NBN, MRE11 and RAD50 mRNA measured by qPCR, normalised as in (B). (E) Levels of NBN, MRE11 protein after 5 days hypoxic (1% O_2_) or normoxic (21% O2) exposure, measured by western-blot. Data shown are ± S.E.M of at least 3 independent experiments. Densitometry quantification for western blots using Image J. * denotes significance (p<0.05) as determined by student t-test.

NBN is responsible for the recruitment and activation of ATM, which then gets activated through auto-phosphorylation at the DNA breakage site^22, 69^ Active ATM phosphorylates downstream targets including the histone variant H2AX and Chk2, ultimately resulting in repair of the break or activation of pro-apoptotic pathways via p53 stabilisation (reviewed by^56^). To understand whether the hypoxia-induced down-regulation of NBN and MRE11 influences ATM activation, we assessed the level of ATM and ATM serine 1981 in hypoxic and normoxic D283-MED cells. Etoposide has the ability to induce replication associated DSB thus triggering ATM activation, measured by an increase in ATM serine 1981 (Figure 5). For cells pre-incubated in 1% O_2_ for 5 days we observed a ~70% reduction in ATM levels compared to the normoxic control. Upon etoposide treatment ATM serine 1981 was ~85% lower in hypoxic cells. This effect of hypoxia on ATM levels was even stronger when cells were exposed to more severe hypoxic conditions (0.1% O_2_) (Figure 5). In contrast, hypoxia had minimal influence on ATM or ATM serine 1981 levels in the U87MG (Figure S3), in line with the previous observation of NBN/MRE11 levels being unaffected by hypoxia in these cells.

**Figure 5:**
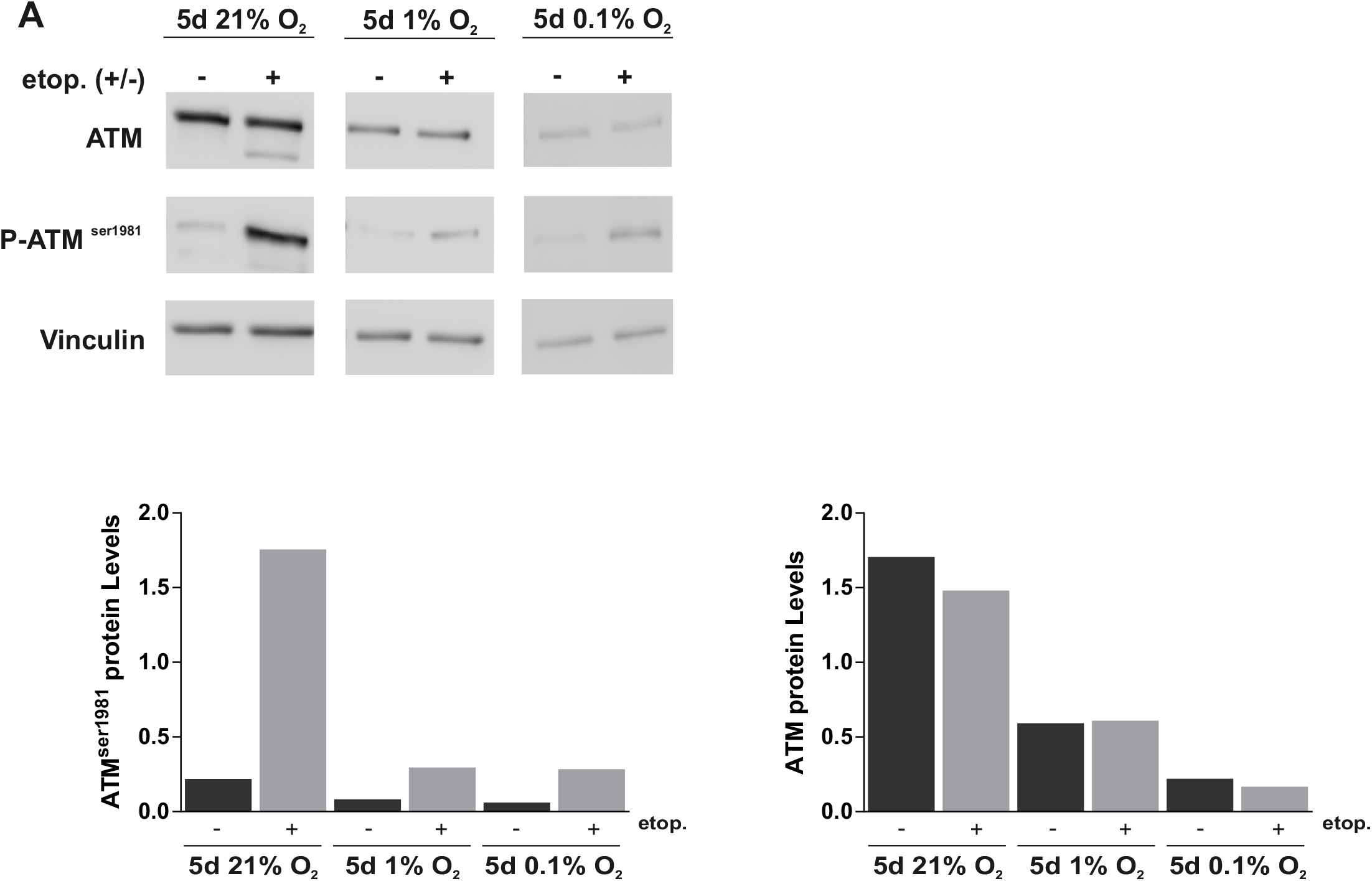
Chronic hypoxia reduces ATM activation after etoposide treatment. D283-MED cells were pre-incubated in 21% O_2_, 1% O_2_ or 0.1% O_2_ prior to treatment with 20μM etoposide for 4 h, or cells were left untreated as a control. Levels of ATM and ATM serine 1981 were determined using 3 independent western blots and densitometry of a representative western blot was measured using Image J.

To ensure that the reduced ATM activation was not a result of decreased etoposide efficacy in hypoxia, we measured the levels of phosphorylated γH2AX, a marker of double strand breaks. In control non-treated cells, there was little γH2AX staining in normoxic and hypoxic cells (Figure 6A), indicating that, as expected, 1% O_2_ is not sufficient to create DSBs in cells. After 15 min of etoposide treatment a similar level of γH2AX levels were observed in normoxia and hypoxia (3.7-fold and 2.9-fold increase, respectively), demonstrating that the efficacy of etoposide in generating DSBs is not affected by the hypoxic conditions alone. This was further confirmed using an alkaline comet assay, which showed that hypoxia had no direct effect on the ability of etoposide to induce DNA strand breaks. Indeed, etoposide induced a clear comet tail in D283-MED cells, and no difference of olive tail moments could be observed between the normoxic and hypoxic cells at any time point (Figure 6B). Together, these experiments demonstrate that etoposide remains fully functional in a low oxygen environment (1% O_2_). These data suggest that as a result of decreased NBN and MRE11 expression in hypoxic conditions there is a clear dampening of further down-stream signalling in the HRR pathway in D283-MED cells.

**Figure 6:**
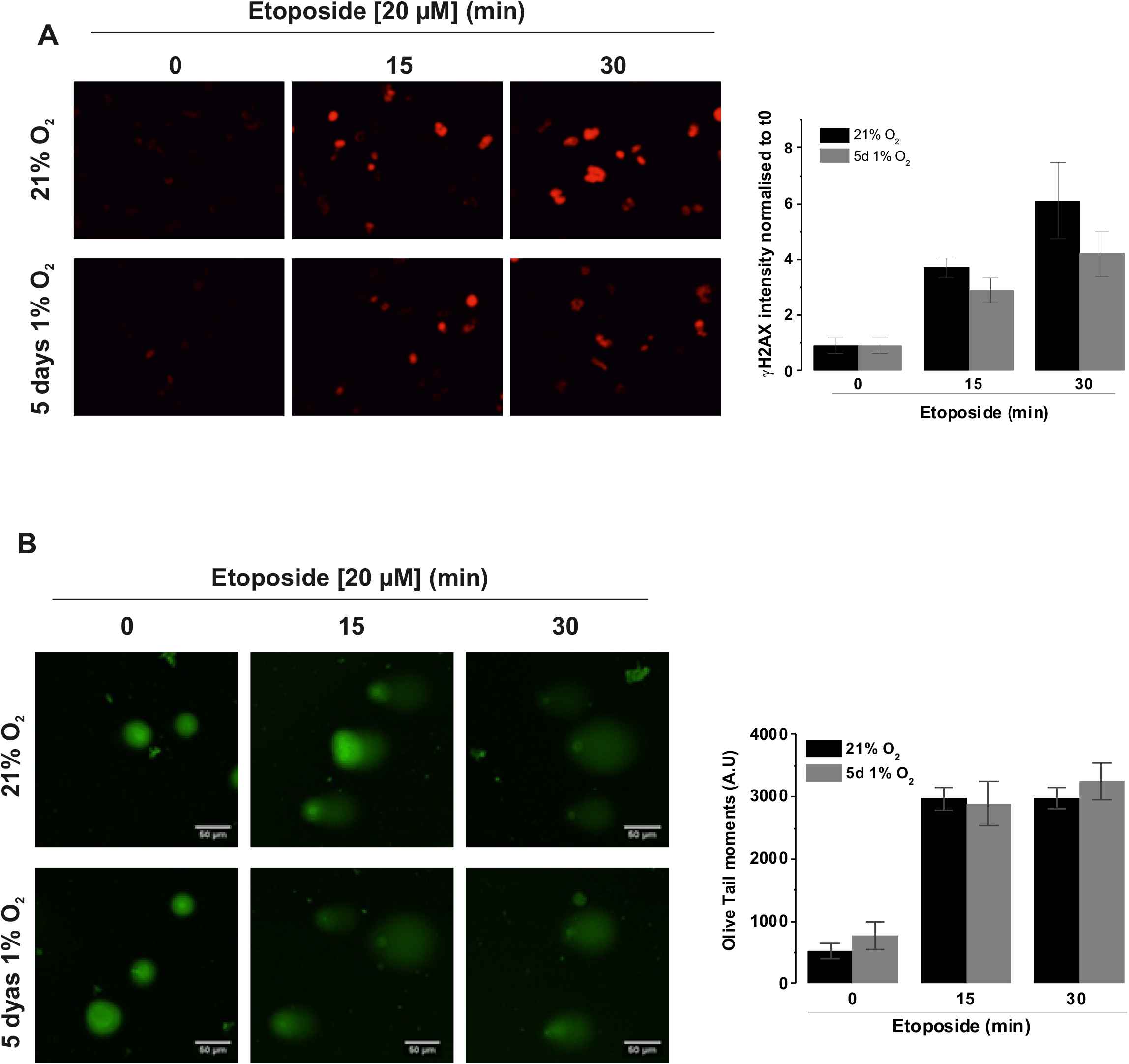
Etoposide efficacy is not affected by hypoxia. D283-MED cells were incubated in 1% O_2_ for 5 days or in 21% O_2_ prior to etoposide (20 μM) treatment for indicated time points. (A) ⁎H2AX bound secondary Cy3 antibodies were detected by immunofluorescence, staining intensity was quantified for individual cells using AQM analysis. Data shown are the mean ± S.E.M of three independent experiments (n >350 cells). One-way ANOVA followed by Bonferroni test was performed (*indicates p<0.05). Images are of a single representative experiment. Quantitative data are represented as a fold change relative to T0. (B) Comet assay (performed according to manufacturer’s protocol). Percentage of ‘Olive Tail moment’ was calculated (see methods section). Results shown are the mean ± S.E.M of 3 independent experiments (n>100cells). One-Way ANOVA followed by Bonferroni test performed (*indicated<0.05).

### p53 activation is reduced in chronic hypoxia

An obvious candidate involved in both cell cycle arrest and apoptosis is p53, a central transcription factor critical for the effectiveness of DNA damaging agents^52^. ATM is directly responsible for p53 stabilisation through phosphorylation^51^. Initially we assessed p53 transcriptional activity in D283-MED by measuring the mRNA levels of three well described p53 target genes: *mdm2* (murine double minute 2), *puma* (p53 upregulated modulator of apoptosis) and *p21*. Hypoxia (5 days at 1% O_2_) did not alter the p53 basal activity, as shown by the similar levels of expression of the target mRNA in all conditions (Figure 7A). As expected, etoposide treatment strongly induced the transcription of *mdm2, puma* and *p21* (~32-, ~5- and ~26-fold, respectively; Figure 7B). However, in cells preconditioned in hypoxia, no increase in *puma* mRNA could be detected and only ~3- and ~2.5-fold increases of *mdm2* and *p21* were observed, which is approximatively 10 times lower than in normoxic cells (Figure 7B). The impairment of etoposide-induced p53 transcriptional activity in hypoxic cells was further confirmed by lower levels of p53 phosphorylation on serine 15 (Figure 7C, D), a marker of p53 activation^52^. Again, basal p53 and p53 serine 15 protein levels were similar in both normoxic and hypoxic cells, yet in hypoxic cells, etoposide failed to fully induce p53 stabilisation with three times lower levels of phosphorylation observed when normalised to the total amount of p53 (Figure 7D). Similar results were obtained for the U87MG glioblastoma cell line (Figure S4A, B), suggesting that, whilst recruiting different mechanisms of resistance, there is a convergence on the activation of p53 protein.

**Figure 7:**
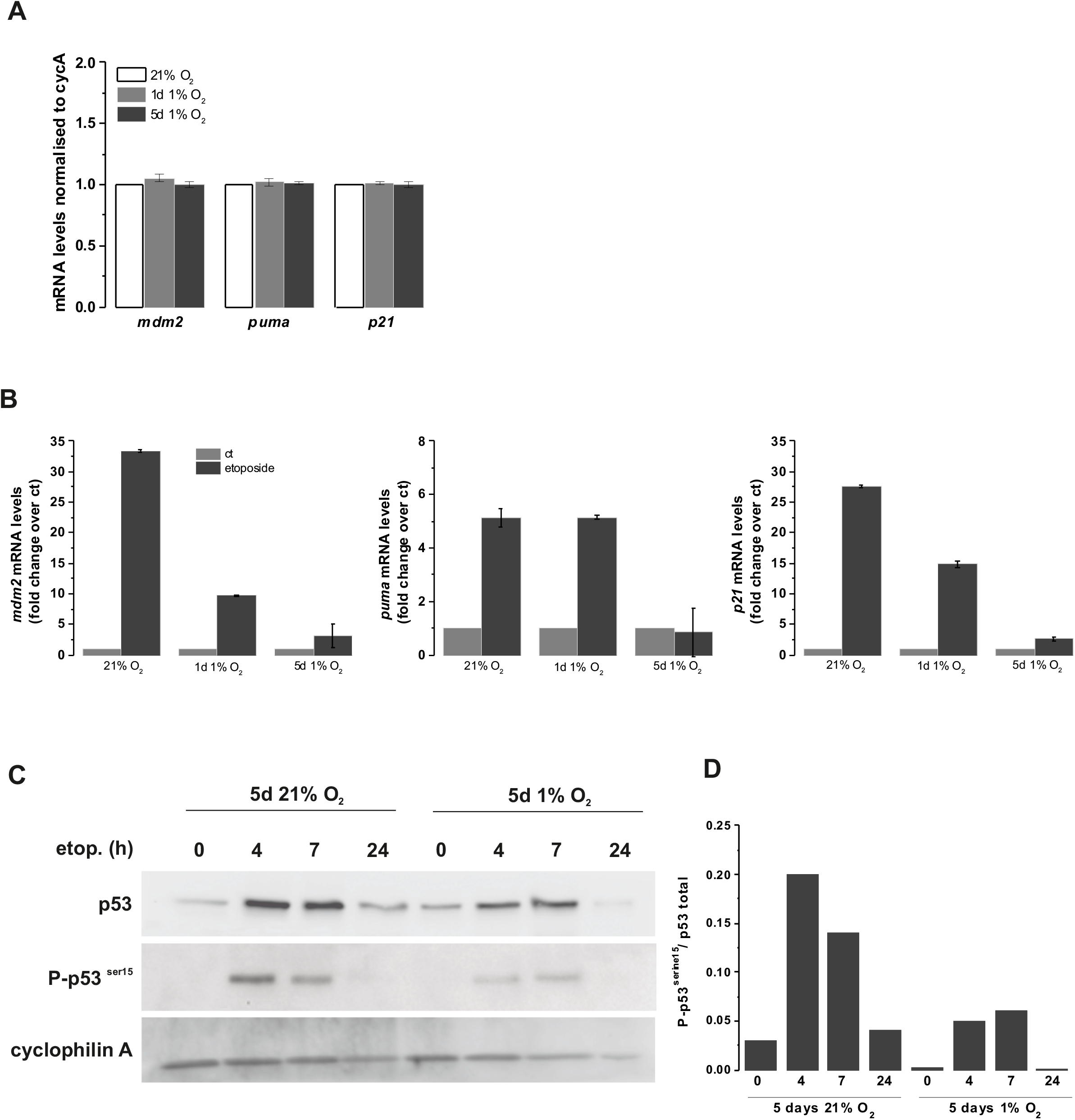
Etoposide induced p53 activity is dampened in chronic hypoxia. D283-MED cells were incubated in 1% O_2_ or 21% O_2_ for 1 day or 5 days, prior to etoposide treatment where indicated. Three p53 target genes, MDM2, PUMA and p21 were assessed by qPCR. (A) Basal levels of p53 target genes in hypoxia without etoposide treatment were measured. The mRNA levels were normalised with the house-keeping gene cyclophilin A. (B) Levels of MDM2, p21 and PUMA mRNA with or without etoposide treatment. Data represented as normalised to housekeeping gene (cyclophilin A) and fold change with respect to the untreated control. Data are representative of three independent experiments, error bars are SD of a single experiment. (C) Total p53 and phosphorylated p53 serine 15 levels assessed by western blot. (D) Densitometry quantification of the band intensity was analysed by ImageJ from 3 independent experiments. Plot represents p53 serine 15 over the p53 total.

In conclusion, we have shown that the chemo- and radio-resistance triggered by a chronic hypoxic exposure can be explained by a broad gene expression remodelling of components of the DNA DSB sensing machinery, subsequently affecting the activation of DNA repair and pro-apoptotic mechanisms. This suggests that reactivation of the signalling pathway could re-sensitise cells to treatment. However, the way cells adapt to the low oxygen levels is diverse with individual cell line or cell-type specific responses. In this study for example, whilst the U87MG cells exposed to chronic hypoxia, showed a similar resistance to treatment and a similar dampened p53 activation to the D283-MED cells, the mechanism of alteration of the DNA DSB sensing and repair pathways were different. Such diversity in the cellular adaptation, makes it difficult to define global clinical strategies for hypoxic cell sensitisation.

## Discussion

Tumour hypoxia has been extensively correlated with acquired resistance to cytotoxic drugs and irradiation, yet the underlying mechanisms are variable between tumour types. The impact of oxygen deprivation during treatment as opposed to the cellular experience prior to treatment has not been untangled so far. Whilst the oxygen enhancement effect, whereby oxygen directly reacts with broken end of DNA preventing their repair, has been implicated in reduced effectiveness of radiotherapy in hypoxia^12, 20^, we here demonstrate that the cellular oxygenation history has a broad impact on cell adaptation and sensitivity to DNA damaging agents. To observe resistance to drug and irradiation a chronic hypoxic preconditioning of several days is necessary, in both medulloblastoma and glioblastoma cells. Although, the prolonged hypoxia exposure triggers different adaptive mechanisms between cell lines, they ultimately converge on the inability to activate the pro-apoptotic p53 protein, thereby mediating cell death and cell-cycle arrest resistance.

### Timing and severity of hypoxic exposure impacts how DNA repair mechanisms are altered

Low oxygen environments such as those experienced within tumours have previously been reported to influence the response to DNA damage and its repair (reviewed in^7 53^). Hypoxia alone has been reported to activate ATM, yH2AX and p53^3, 26, 67^, however, this was not the case in our hypoxic conditions (1% O_2_), which were much milder than those used in these previous studies (0-0.2% O_2_). Additionally, severe hypoxia has been reported to reduce HRR capacity by the down-regulation of key repair proteins such as RAD51 and BRCA1^5, 6^^11^. Work by To *et al*. identified that moderate hypoxia (1% O_2_) resulted in transcriptional downregulation of NBN through HIF-1α binding to the NBN promoter thereby displacing Myc^64^, supporting our own findings (Figure 4). Such impact of chronic hypoxia on HRR was reported to result in increased sensitivity to DNA damaging substances such as crosslinking agents (Cisplatin, Mytomycin C) and irradiation^11, 34^. Our data suggest that hypoxia-induced downregulation of the MRN complex results in resistance to DSB inducing agents by impairing signalling events downstream of the MRN recruitment hence resulting in reduced p53 activity and apoptosis.

The differing O_2_ levels experienced by cells, are varied in terms of localisation within the solid tumour and duration of the hypoxic episodes^65^, including cycling hypoxia^21^. *In vitro* experiments, trying to recreate such variety of hypoxic severity and duration have led to conflicting results rendering studies hard to compare^2^. We identified that hypoxia-induced resistance to etoposide required chronic (5 day), rather than acute (24 h), hypoxia. Previous studies investigating hypoxia and drug resistance primarily utilised acute (6-48 h) time points. For example, resistance to cisplatin and doxorubicin induced by hypoxia in non-small cell lung cancer was due to incubation in 0.5% O_2_ for only 16 h^59^. Our work highlights the necessity to explore both chronic and acute exposure times to ensure adequate observation of any potential hypoxia induced resistance in cancer models. When considering the impact of hypoxia duration on DNA repair mechanisms, short exposures (minutes to hours) are thought to regulate repair proteins through post-translational modifications and changes to translation efficiency, whereas longer exposure (days-weeks) results in alteration to transcription, as well as epigenetic modifications^53^. The reported activation of numerous DNA repair proteins by hypoxia, such as ATM and p53, as well as the reduced capacity of HRR was primarily observed after acute and severe hypoxic exposure (16-72 h)^3, 5, 6, 11, 26, 34^ Our gene expression analysis identified a significant proportion of genes involved in DNA DSB repair that were only regulated at later hypoxic time points (64-96 h). This highlights the need to extend hypoxia incubation periods for DNA repair related studies. Thus far the impact of more chronic moderate hypoxia on DNA repair mechanisms and the biological and clinical implications may have been missed.

### Heterogeneity of the cellular response to hypoxia between cell types

We observed a strong hypoxia-induced chemo- and radio-resistance in the D283-MED cell line (Figure 1), however the MEB-Med8A cells did not show the same extent of resistance. This differing response might be due to the differences in the molecular signature of these two cell lines. MEB-Med8A is a Group 3 medulloblastoma cell line, with strong associations with classical Group 3 characteristics, such as *myc* amplification and *pvt1-myc* fusion^35, 44^, whereas the grouping affiliation of D283-MED is more controversial, with a lack of *myc* amplification pointing towards Group4^4, 54^. Additionally, MEB-Med8A has a truncated form of p53, which impacts on the effectiveness of response to DNA damaging agents^39^. Such distinct genetic backgrounds are likely to explain their differing response to hypoxia, reinforcing the view that the genetic make-up of a tumour is a key factor to consider when selecting therapeutic regimens. In addition to the resistance observed in D283-MED cells, similar effects of hypoxic preconditioning were observed in U87MG cells, a classic glioblastoma cell line, in agreement with previous reports^15, 30^. However, in the U87MG cells, hypoxia did not influence the level of the MRN complex or ATM activation (Figure S3) as it did in the D283-MED cells, demonstrating again the heterogeneity in the cellular response to hypoxia, beyond HIF activation. Additionally, we did not observe other reported mechanisms of hypoxia-induced resistance such as the induction of MDR genes^13^ in the D283-MED cells.

The complex and varied response to hypoxia is also apparent when comparing *in vitro* cell line models to *in vivo* organisms and pre-clinical models. In this case, our results obtained *in vitro* mimicked the observations made in hypoxic tumours *in vivo*, in terms of resistance to cancer treatments. Moreover, hypoxia related changes to DNA repair proteins have been translated to an *in vivo* setting. For example, active phospho-ATM is co-localised with the hypoxic marker CAIX in tumour xenografts^45^ and RAD51 down-regulation was observed in cervical and prostate cancer xenografts^5^ as well as in a glioma model^37^.

In a clinical perspective, the development of HIF inhibitors targeting either HIF mRNA transcription, translation, or HIF stabilisation^38^ might help tackling the treatment resistance problem by re-sensitising hypoxic tumours. In the U251 glioblastoma cells, campothecins (CPTs) analogues can inhibit the accumulation of HIF1a protein^49^ However, individual tumours react differently to hypoxic episodes, as highlighted by our own data, which exemplifies the need to understand how hypoxia impacts tumour cell biology on an individual tumour basis leading to a more personalised clinical approach.

## Material and Methods

### Reagents

Etoposide (E1383) was from Sigma. Tissue cell culture mediums were supplied by Gibco Life Technologies and foetal calf serum from Harlan Seralab (UK). Cyclophilin A (Ab3563), anti-mouse (Ab6808), MRE11 (ab214), ATM (ab3240), ATM serine 1981 (ab81292) and Vinculin (ab129002) antibodies were from Abcam. p53 BC-12 (sc-126) and NBN (sc-8589) antibodies were from Santa Cruz. p53 serine 15 (PC386) antibody was from CalBiochem. Anti-rabbit (7074) antibody was from Cell Signalling. Cy3-anti mouse antibody was from sigma (C2181).

### Cell culture

D283-MED and U87MG were purchased from ATCC. MEB-Med8A cells were kindly provided by Prof T. Pietsch (University of Bonn, Germany). D283-MED were maintained in modified Eagle’s medium (MEM) with 10% FCS, 1% non-essential amino acid and 1% sodium pyruvate. MEB-Med8A cells were maintained in Dulbecco’s MEM (DMEM) with 10% FCS. U87MG cells were maintained MEM with 10% FCS and 1% sodium pyruvate. Cells were typically cultured and maintained at 37°C and 5% CO2 in a humidified incubator. For hypoxic conditions (1% O_2_) cells were manipulated and incubated in a hypoxic workstation (Don Whitley Scientific, UK).

### Quantitative Real time PCR (qPCR)

RNA was extracted with RNeasy Mini kit (Qiagen, Germany). 1μg of RNA were used for cDNA synthesis using SuperScript^®^ VILO kit according to the manufacturer’s protocol (Invitrogen). Quantitative qPCR was performed as described in 50 Primers sequences were as follows: Cyclophilin A: Forward: GCTTTGGGTCCAGGAATGG; Reverse: GTTGTCCACAGTCAGCAATGGT; MDM2: Forward: GCAAAT GT GCAAT ACCAACA; Reverse: CTTT GGT CT AACC AGGGT CT C; PUMA: CCTGGAGGGT CCT GT ACAAT; Reverse: C ACCT AATT GGGCT CAT CT; p21: G ACT CT C AGGGT CG AAAACG; Reverse: T AGGGCTT CCT CTT CCAGAA; NBN AG AATT GGCTTTT CCCG AACT; Reverse: CAAGAAGAGCAT GC AACCA. MRE11 Forward: T CAGT CAAGCT CCT CT GGG A; Reverse: AGT CC AGCAGT GGG AATTT CT; RAD50: TGCTTGTTGAACAGGGTCGT; Reverse: TCACTGAATGGTCCACGCTC. Fold change was calculated based on the threshold of amplification cycle for each reaction using the 2-CT method^46^, where target genes were normalised to cyclophilin A and the control.

### Microarray Analysis

D283-MED cells were cultured in 10cm dishes. Reverse transcriptome amplification of extracted RNA was conducted using Transplex^®^ Whole Transcriptome Amplification kit (Sigma) from 3 different replicate per time point. NimbleGen 12x135k format array slide was utilised for the microarray experiment, whereby each transcript was represented by 3 probes. Statistical analysis of data was conducted using Matlab (script written by Damon Daniels, University of Manchester), with clustering based on *k*-means. All microarray raw and normalised data are available on NCBI: <u>http://www.ncbi.nlm.nih.gov/geo/query/acc.cgi?acc=GSE106959</u>)

### MTS assays

For irradiation treatment, cells were treated in 35mm dishes with 30-80 Gy x-ray irradiation in a CellRad Faxitron (130-150 kV, 5 mA), before seeding into a 96 well culture plates. Etoposide treatment was conducted directly in pre-seeded 96 well plates, with treatment of 20μM-100μM etoposide. CellTiter 96^®^Aqueous One Solution (Promega) was added to wells and incubated for 2-4 h at 37°C at the end of each treatment time point. Measurements were obtained with a plate reader at 492 nm (Multiskan, Thermo Scientific).

ViaCount assays: D283-MED cells were cultured on 6 well plates, 24 h prior drug treatments. The cells were treated with etoposide (20 μM) or left untreated (control). At the end each time point, cells were prepared according manufacturer protocols for FACS analysis. Samples were ran using a ViaCount analysis on a GUAVA flow cytometry (Guava EasyCyte Plus). Percentages of viable cells were measured using the Guava ViaCount software.

Cell cycle analysis: D283-MED cells were cultured on 6 well plates in either normoxia or hypoxia, with or without etoposide treatment. The preconditioned cells were centrifuged and resuspend in HBSS (200 μl) and stained with 25 μl of propidium iodide [10 μg/ml]. FACS analysis was performed using GUAVA flowcytometry (Guava EasyCyte Plus). Percentage of cells in each cell cycle phase was calculated using Guaava Express Pro software.

### Immunostaining

D283-MED were plated on polyornithine [100 μg/ml] pre-treated coverslips 16 hours prior to treatment. Cells were fixed with paraformaldehyde (4%), and washed in PBS. PBS containing 1% BSA and 0.1% Triton-X was used for blocking and permeabilisation of cells, followed by primary antibody incubation with γH2AX for 16 h, and secondary antibody incubation for 30 min. Cells were then stained with DAPI (Hoechst, 1 μl/ml in PBS) for 5min. Coverslips were mounted onto glass slides using Dako mounting medium. Images were obtained using Leica DM2500 microscope (Leica) and quantified using AQM Advance 6 software (Kinetic Imaging Ltd, UK).

### Western Blotting

Conducted as described in^50^. In brief, 20-40 μg of protein was resolved on a 10% SDS-PAGE gel and transferred to nitrocelloulose membrane (0.2 μM) before incubation in primary and secondary antibody. Amersham ECL Prime Western Blotting Detection Reagent (GE Healthcare) was used for development of signal, and a G:BOX gel imaging system (Syngene, UK) used for detection. Western blot quantification was conducted with ImageJ 1.45s open source software (National Institutes of Health, USA).

### Comet assay

D283-MED cells were cultures on a 12 well plate directly before drug treatments. Cells were collected by centrifugation and re-suspended in 1 ml ice-cold PBS. The samples were diluted with pre-warmed agarose and loaded on the slide provided by the OxiSelectTM comet assay kit. Subsequently, the Comet assay procedures were performed as per manufacturer’s protocol, followed by alkaline electrophoresis. Vista dye was added immediately after electrophoresis for DNA staining. Slides were then imaged using Leica

### DM2500 microscope

Images were analysed and quantified using the ImageJ 1.45s open source software (National Institutes of Health, USA).

### Statistical analysis

Statistical significance test was performed using OriginPro 8.6.0 (OriginLab Corporation, USA).

## Acknowledgments

C.F. and S.C. were funded by Alder Hey Children Hospital charitable funds. Immunostaining imaging was performed in the Liverpool Centre for Cell Imaging using equipment funded by the MRC (MR/K015931/1). We thank Dr Nigel Jones and Prof Dave Fernig (University of Liverpool) for critical reading of the manuscript. We thank Dr Jason Parson (University of Liverpool) for access to the CellRad Faxitron irradiator and Dr Helen Wright (University of Liverpool) for her support with cell cycle analysis. We thank Damon Daniels (University of Manchester) for his help with micro-array analysis and Matlab coding

## Supplemental Figures

**Figure S1: Hypoxia alone does not affect the cell cycle of D283-MED cells**. (A) D283 cells were pre-incubated in 1% O_2_ for 5 days (grey line) or left in normoxia 21% O_2_ (black line). Cells were treated with etoposide (20 μM) for the indicated time. Cell survival was assessed using ViaCount assay. Results are expressed as percentage cell survival relative to the untreated control. Data shown are the mean ± S.E.M of three independent experiments. One-Way ANOVA followed by a Bonferroni test was performed (* indicates p < 0.05). (B) Cells were incubated in normoxia or 5 days hypoxia and treated with etoposide (1 μM) for 30 hours. Cell cycle profile obtained from the Guava software. Data shown are representative of 3 independent experiments. Quantitative data showing the percentage of cells in each phase, value is calculated by dividing the number of cells in each phase by the total number of cells for that sample. Data shown are the mean ± S.E.M of 3 independent experiments. (C) Cell cycle profiles obtained from flow cytometry. D283 cells were cultured in normoxia or hypoxia (1% O_2_) for indicated times. Cell cycle profile quantification showing percentage of cells in each cell cycle phase. Data representative of 3 independent experiments, error bars shown are experimental errors of 2 replicates.

**Figure S2: *k*-means clustering of regulated transcripts**. Significantly regulated transcripts from microarray data clustered into one of 16 groups using k-means. Transcripts with similar expression level are grouped together in the same cluster.

**Figure S3: Expression of the MRN complex and ATM activation is not affected by chronic hypoxia in U87MG cells**. U87MG cells were pre-incubated in 21% O_2_, 1% O_2_ or 0.1% O_2_ for 5 days prior to 4hr 100μM etoposide treatment where indicated. (A) mRNA levels of NBN, MRE11, RAD50 were determined by qPCR, normalised to the housekeeping gene cyclophillin A and shown as fold change relative to normoxic control. Data represent mean ± S.E.M of at least 3 independent experiments. * denotes significance (p<0.05) as determined by student t-test. (B) Levels of ATM and ATM serine 1981 determined using western blotting and densitometry of a representative western blot measured using ImageJ.

**Figure S4: Etoposide induced p53 activity is dampened by chronic hypoxia in U87MG cells**. U87MG cells were incubated in 1% O_2_ or 21% O_2_ for 5 days, prior to etoposide treatment where indicated. Three p53 target genes, MDM2, PUMA and p21 were assessed by qPCR. (A) Levels of MDM2, p21 and PUMA mRNA with or without etoposide treatment. Data represented as normalised to housekeeping gene (cyclophilin A) and fold change with respect to the untreated control. Data are representative of a single experiment. (B) Total p53 and phosphorylated p53 serine 15 levels assessed by western blot. Densitometry quantification of the band intensity was analysed by ImageJ of a single experiment. Plot represents p53 serine 15 normalised over the p53 total.

